# Adaptive History Biases Result from Confidence-weighted Accumulation of Past Choices

**DOI:** 10.1101/172049

**Authors:** Anke Braun, Anne E. Urai, Tobias H. Donner

## Abstract

Perceptual decision-making is biased by previous events, including the history of preceding choices: Observers tend to repeat (or alternate) their judgments of the sensory environment more often than expected by chance. Computational models postulate that these so-called choice history biases result from the accumulation of internal decision signals across trials. Here, we provide psychophysical evidence for such a mechanism and its adaptive utility. Male and female human observers performed different variants of a challenging visual motion discrimination task near psychophysical threshold. In a first experiment, we decoupled categorical perceptual choices and motor responses on a trial-by-trial basis. Choice history bias was explained by previous perceptual choices, not motor responses, highlighting the importance of internal decision signals in action-independent formats. In a second experiment, observers performed the task in stimulus environments containing different levels of autocorrelation and providing no external feedback about choice correctness. Despite performing under overall high levels of uncertainty, observers adjusted both the strength and the sign of their choice history biases to these environments. When stimulus sequences were dominated by either repetitions or alternations, the individual degree of this adjustment of history bias was about as good a predictor of individual performance as individual perceptual sensitivity. The history bias adjustment scaled with two proxies for observers’ confidence about their previous choices (accuracy and reaction time). Taken together, our results are consistent with the idea that action-independent, confidence-modulated decision variables are accumulated across choices in a flexible manner that depends on decision-makers’ model of their environment.

**Significance statement:** Decisions based on sensory input are often influenced by the history of one’s preceding choices, manifesting as a bias to systematically repeat (or alternate) choices. We here provide support for the idea that such choice history biases arise from the context-dependent accumulation of a quantity referred to as the decision variable: the variable’s sign dictates the choice and its magnitude the confidence about choice correctness. We show that choices are accumulated in an action-independent format and a context-dependent manner, weighted by the confidence about their correctness. This confidence-weighted accumulation of choices enables decision-makers to flexibly adjust their behavior to different sensory environments. The bias adjustment can be as important for optimizing performance as one’s sensitivity to the momentary sensory input.

## Introduction

It has been known for almost a century that people’s judgments of sensory stimuli do not only depend on the current sensory input, but also on their preceding choices (Fernberger, 1920). Several studies have found that humans and other species repeat (or alternate) their perceptual judgments more often than expected by chance (Gold et al., 2008; Busse et al., 2011; de Lange et al., 2013; Akaishi et al., 2014; Fischer and Whitney, 2014; Fründ et al., 2014; Abrahamyan et al., 2016; Pape and Siegel, 2016; St. John-Saaltink et al., 2016; Fritsche et al., 2017; Hwang et al., 2017; Urai et al., 2017). Such choice history biases occur also in other domains of decision-making (Leopold et al., 2002; Allefeld et al., 2013; Padoa-Schioppa, 2013).

Computational models posit that choice history biases result from the temporal accumulation of signals from past decisions (Yu and Cohen, 2009; Glaze et al., 2015; Bonaiuto et al., 2016). Such a mechanism may serve to continuously update the decision-makers’ prior belief about the upcoming stimulus category and adjust their choice behavior to structured environments (Yu and Cohen, 2009; Glaze et al., 2015). In laboratory perceptual tasks, stimulus sequences are typically uncorrelated by design, so that across-trial accumulation degrades performance (Abrahamyan et al., 2016). By contrast, when stimulus sequences exhibit auto-correlations (Goldfarb et al., 2012; Glaze et al., 2015; Abrahamyan et al., 2016; Kim et al., 2017), history biases should improve performance, provided that the accumulation is context-dependent. Specifically, the accumulation should switch sign between environments dominated by either stability or change (Glaze et al, 2015).

Perceptual decisions often have to be made under uncertainty due to weak or ambiguous evidence. This uncertainty (or its complement: confidence) might be important for controlling behavior under conditions, in which the decision-maker receives no immediate external feedback. Indeed, fluctuations of confidence play a key role in a normative model, which postulates the accumulation of the internal decision variable over time (Glaze et al, 2015). The decision variable is the basis of both the categorical choice (Bogacz et al., 2006; Gold and Shadlen, 2007) as well as the confidence about its correctness (Kepecs et al, 2008). Correlates of the decision variable are distributed across many brain regions (Gold and Shadlen, 2007 Siegel et al., 2011; Brody and Hanks, 2016) and expressed as motor plans (Gold and Shadlen, 2007; Donner et al., 2009; de Lange et al., 2013) or in action-independent formats (Bennur and Gold, 2011; Hebart et al., 2012; O’Connell et al., 2012; Hebart et al., 2016). These decision-related neural signals also reflect the graded confidence about the choice (Kiani and Shadlen, 2009; Hebart et al., 2016).

Our current study addressed three questions. First, do choice history biases originate from signals in motor or action-independent formats? Second, can these signals be accumulated in a sufficiently flexible manner, so as to adjust history biases to repetitive as well as alternating environments? Third, is the strength of such bias adjustment scaled by confidence? We modeled human choice behavior under experimental manipulations tailored to answering these questions.

## Materials and Methods

### Participants

We analyzed data from 28 participants and two experiments (referred to as Experiment 1 and 2) in total. All participants gave their written informed consent.

#### Experiment 1

Six healthy participants (2 male and 4 female, mean age: 25; range: 22–29 years) took part in the experiment, which was approved by the ethics committee of the Department of Psychology of the University of Amsterdam (reference number 2011-OP-1588).

#### Experiment 2

26 healthy participants (11 male and 15 female, mean age: 26, range: 20–36) took part in the experiment, which was approved by the local ethical review board (Ärztekammer Hamburg, reference number PV4714). Four participants were excluded from the data analysis, so that 22 participants remained for the data analysis. Three of the excluded participants did not complete all sessions and one exhibited substantially worse performance than the rest of the group (64 percent correct overall, 63 percent correct for the easiest three motion coherence levels).

### Experimental design

The data from both experiments allowed for quantifying choice history biases during a random dot motion discrimination (up vs. down) task. We used large random dot motion patterns in both experiments, so as to minimize stochastic fluctuations in the effective motion energy across trials (Urai et al., 2017).

#### Experiment 1

The following description summarizes the aspects of the experimental design that were most important to the current paper; a comprehensive description can be found in (Tsetsos et al., 2015). Random dot kinematograms (Figure 1A) were composed of 785 (average) white dots on a black screen. The dots were moving within a circular aperture of 9.1° radius. A red fixation cross of 0.4°×0.4° was centered in the middle of the circle. The dot density was 12.07 dots per deg^2^. The population of dots was split into “signal dots” and “noise dots”. The signal dots moved either upwards or downwards with a velocity of 2.6°/s. The noise dots changed position randomly from frame to frame. The percentage of signal dots defined the motion coherence, a measure of motion strength. On each trial, three different sequences of dot motion (at the same coherence and direction) were presented in an interleaved fashion, making the effective speed of signal dots 0.87 °/s. One of six different levels of motion coherence, (0.05, 1.26, 3.15, 7.92, 19.91 and 50%) and one of six different viewing durations (150, 300, 600, 1200, 2400, and 4800 ms) were chosen randomly, under the constraint that they occurred equally often within a block of 144 trials. Stimuli were presented on a 22-inch CRT monitor with a resolution of 800×600 pixel and a frame rate of 100 Hz at a viewing distance of 68 cm. The participants were instructed to maintain their gaze on the red cross throughout the trial and judge the net motion direction. The motion viewing interval was followed by a variable delay (uniform distribution ranging from 200 to 400 ms), after which the observers had to report their choice by pressing one of two buttons on a computer keyboard, with either the left or the right index finger. Participants received auditory feedback after incorrect responses (a 1000 Hz tone of 100 ms). Perceptual choices (‘up’ vs. ‘down’ motion direction) were decoupled from motor responses (left vs. right button press) by varying their mapping from trial to trial. This mapping was instructed before motion viewing in one condition (‘Pre’ condition) and after motion viewing in the other (‘Post’ condition), by means of a visual cue that presented each direction (as an arrow) on the left or right side (i.e. two possible mappings). This mapping cue was randomly selected on each trial. Conditions alternated across blocks. Observer 1-5 participated in both conditions. Observer 6 participated only in the Post condition. The analyses of participants 1-5 presented here were collapsed across both conditions. We obtained the same pattern of results when analyzing the data from both conditions separately (data not shown).

**Figure 1.**
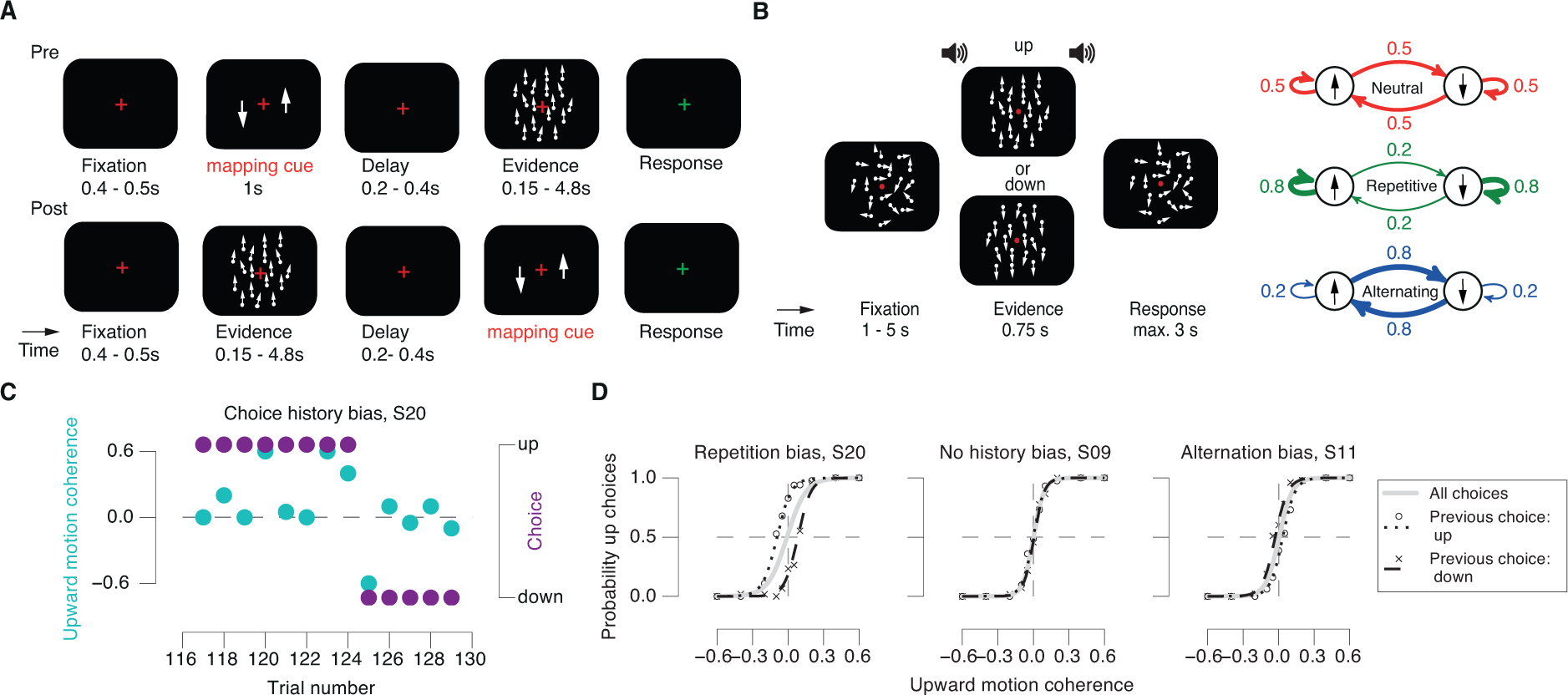
Quantifying choice history bias and behavioral task. (**A**), (**B**) Behavioral tasks. Observers judged the net direction (up vs. down) of a dynamic random dot pattern of variable direction and coherence. (**A**) Experiment 1, decoupling choice and motor response. After a blank fixation interval, a choice-response mapping cue was shown before (Pre) or after the presentation (Post) of the motion stimulus, which also varied in duration. Observers responded after dot motion offset in the Pre-condition and after mapping cue offset in the Post-condition. Auditory feedback was provided after incorrect responses. (**B**) Experiment 2, manipulating stimulus repetition probabilities. Left: Random dot motion and fixation cross were shown throughout the trial. A beep indicated the onset of the evidence interval, which contained some level of coherent motion (0% on some trials). A second beep indicated the evidence offset and start of the response interval (deadline: 3 s). Right: Three repetition probabilities between motion directions across trials yielded three environmental conditions: Neutral (repetition probability of 0.5), Repetitive (repetition probability of 0.8) and Alternating (repetition probability of 0.2). (**C**) Signed motion coherence levels (cyan) and categorical choices (purple) from a sequence of 15 trials recorded in Neutral in Experiment 2. Positive values of stimulus intensity correspond to upward motion and negative ones to downward motion. (**D**) Psychometric functions conditioned on previous choice in Neutral exhibit history biases in three example participants. See main text for details.

#### Experiment 2

To test for the adaptability of choice history biases, we manipulated the sequential stimulus statistics between experimental sessions, to make people perform the task in ‘Repetitive’, ‘Neutral’ (no sequential dependence), or ‘Alternating’ environments (Figure 1B). Stimuli, task, and procedure for Experiment 2 were identical to Experiment 1, with the following exceptions. The circle within which the dots were moving had an outer radius of 12° and an inner radius of 2°. The density of dots was 1.7 dots/deg^2^ and each dot had a diameter of 0.2°. The dots moved with a velocity of 11.5°/s. Signal dots had a maximum lifetime of 6 frames. We used the following coherence levels: 0, 5, 10, 20, 40 and 60% (equally many trials per coherence level). A red bulls-eye fixation target at the center of the screen as well as randomly moving dots (0% coherence) were presented throughout each block. The first trial of each block started with a baseline interval of 5 s. A beep (duration: 50 ms, sine wave at 440 Hz) indicated the onset of the evidence interval with variable coherence levels and directions (see above) after a fixed duration of 0.75 s. A second beep indicated the offset of the evidence interval and prompted the observers’ response. Observers reported their perceptual choices by pressing one of two keyboard buttons, with the index finger of the left or right hand. After button press or a response deadline of 3 s, the inter-trial interval started. Inter-trial intervals were uniformly distributed between 1 and 5 s. Observers received auditory feedback during the training sessions, but no feedback during the subsequent six sessions of the main experiment. The motion viewing duration of 0.75 s was selected because previous work in monkeys (Kiani et al., 2008) and humans (Tsetsos et al., 2015) found little integration of motion information beyond that duration. We used a fixed mapping between choices and motor responses, whereby the two possible mappings (right-hand button for up, left-hand for down, or vice versa) were counterbalanced across participants. Experiment 2 consisted of seven sessions per participant (one for training and six main sessions), whereby each session was divided into 10 blocks of 60 trials.

Critically, the transition probabilities between the two alternative stimulus categories (i.e. up vs. down regardless of coherence) over trials were manipulated across experimental sessions (Figure 1B, right). Specifically, the probability of a repetition was defined as

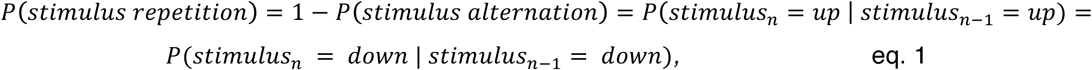

whereby *n* indexes trials. The repetition probability was held constant within each session, but varied across the main experimental sessions between the following values: 0.5 in the ‘Neutral’ condition, 0.8 in the ‘Repetitive’ condition, and 0.2 in the ‘Alternating’ condition. The Neutral condition allowed for quantifying observers’ intrinsic choice history bias, which we used as a baseline for quantifying their adjustment to the biased sequential statistics of the Repetitive and Alternating conditions.

During the training session, the motion direction on each trial was chosen randomly and independently. All participants started with the Neutral condition in session 1 of the main experiment, which was repeated in session 4. Half of the participants then performed the Repetitive condition in sessions 2 and 5 and the Alternating condition in sessions 3 and 6 and conversely for the other half of participants.

Observers were instructed to maintain stable fixation and perform the motion discrimination task as accurately as possible. They were informed that the sequential statistics of the stimulus identities would change from session to session, but stay constant within each session. To this end, we told them that the stimulus sequences could be ‘as if produced by a coin flip’ (Neutral), ‘more likely repeating than alternating’ (Repetitive), or ‘more likely alternating than repeating’ (Alternating). Observers were not informed about (i) the order of these conditions, (ii) the exact transition probabilities, (iii) the use of this information for optimizing their behavioral performance.

### Modeling choice history bias

We used logistic regression to model observers’ choice history biases under the different experimental conditions. The basic approach consisted of adding a linear combination of different components of trial history (which depended on the experiment), as a bias term to a logistic function model of the choice probability (Fründ et al., 2014; Urai et al., 2017). We here used a variant that quantified the relative contributions of previous stimuli, choices, and (for Experiment 1) motor responses.

#### *Basic choice model* using *psychometric function fit*

The probability of making one of the two choices *r_t_* = 1 (*r_t_* = 1 for ‘choice up’, *r_t_* = −1 for ‘choice down’) on trial t, given the signed stimulus intensity 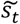 (i.e., motion coherence times stimulus category) (up or down, coded as 1 and −1) was described by:

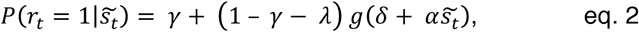

where *γ* and *λ* were the lapse rates for the choices *r_t_* = 1 and *r_t_* = −1, and 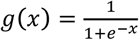 was the logistic function. The bias term *δ*, the offset of the psychometric function, described the overall bias for one specific choice. *α* was the slope of the stimulus-dependent part of the psychometric function, quantifying perceptual sensitivity.

For visualizing the effect of previous on current choice (Figure 1D), we separated the trials from Neutral into two subsets, conditioned on the choice from the previous trial, and fitted the psychometric function separately to the observed proportion of upward choices in both subsets. Results from three example observers are shown in Figure 1D and discussed in Results.

#### Modeling the contributions of past stimuli, choices, and motor responses to current choice bias

We estimated the contribution of the previous seven stimulus categories and choices by adding a history-dependent bias term to the argument of the logistic function (Fründ et al., 2014):

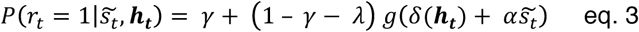

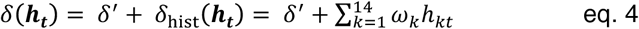

The history bias 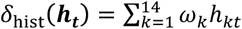 consisted of the sum of the preceding seven responses *r*_*t*−1_ to *r*_*t*−7_ and the preceding seven stimulus categories *z*_*t*−7_ to *z*_*t*−7_, each multiplied with a weighting factor *ω_Λ_*. The vector ***h***_*t*_ was written as:

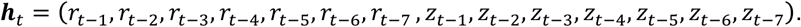

All terms in ***h***_*t*_ were coded as −1 or 1, with the exception of terms coding for stimuli with zero coherence, which were set to zero. The weighting factors *ω_t_* thus modeled the influence of each of the seven preceding responses and stimulus categories on the current choice. Positive values of *ω_t_* indicated a bias to repeat the choice or stimulus category at the corresponding lag, and negative values of *ω_t_* indicated a tendency to alternate. In this and all subsequent analyses, the parameters of the logistic regression model were fit by maximizing the log-likelihood 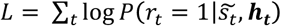 using an expectation maximization algorithm (Fründ et al., 2014).

In Experiment 1, perceptual choices and motor responses were further decoupled through a mapping that varied from trial to trial. Thus, we could independently estimate the relative contribution of previous choices and motor responses to the current choice bias. We added the last seven choices *c_t−1_, c_t−2_, c_t−3_, c_t−4_, c_t−5_, c_t−6_, c_t−7_*, each one multiplied with a separate set of history weights *ω′_k_*, to the history bias term *δ_hist_*(***h_t_, c_t_***).

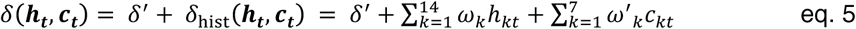

Experiment 1 contained not only trial-to-trial variations in motion direction and coherence, but also in the duration of the dot motion stimulus. To assess the effect of this manipulation, we first fitted the psychometric functions separately for each of the six different motion-viewing durations (on the current trial) and compared the resulting weights within each observer. The viewing duration had only negligible impact on the history weights (data not shown), indicating that the history contributions were invariant across viewing durations. Consequently, we fitted the psychometric functions to the data from all trials, and analyzed the history weights, irrespective of viewing duration.

Experiment 1 also contained two conditions (Pre and Post), in which observers were instructed about the required mapping between choice and response either before or after the presentation of the sensory evidence. The analyses presented in Results collapsed across both conditions, but we also verified that there were no differences between these conditions when analyzing them separately (data not shown).

#### Modeling the contributions of past correct (and incorrect) choices to current bias

The weights for previous correct and incorrect choices were estimated by re-combining the weights for previous stimuli and choices estimated by means of equations 3 and 4. Specifically, the weights for correct choices were computed as the sum of choice and stimulus weights and the weights for incorrect choices were computed as the difference between choice and stimulus weights (Fründ et al., 2014). Please note that this is equivalent to fitting a regression model with predictors encoding correct or incorrect choices, along with the chosen category (Busse et al, 2011; Abrahamyan et al, 2016).

#### Modeling the contribution of past decision confidence to current bias

We used a model of statistical decision confidence based on signal detection theory (Kepecs et al. 2008; Sanders et al. 2016; Urai et al. 2017) in order to define two behavioral proxies of confidence that could be used in the present study. The model assumes that choices are made based on an internal decision variable (*dv*), which is computed as a transformation of sensory input, corrupted by noise. A choice is made by comparing *dv* to a criterion *c.* Confidence is a function of the distance between *dv* and *c.* When *dv* is far from c, the choice is likely to be correct; the probability of the choice being correct approaches chance as *dv* approaches c. Specifically, *confidence*, = *f*(|*dv_i_* − *c*|) where 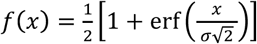 is a monotonic sigmoid function that maps the distance metric×on the probability of making a correct choice (Kepecs et al, 2008; Lak et al. 2014). The model predicts that confidence (i) is larger on correct than on error trials, and (ii) scales oppositely as a function of stimulus strength for correct and error trials. We used two behavioral proxies of the so-defined confidence to investigate its impact on the adjustment of choice history biases: (i) Choice accuracy, with correct choices being associated with larger confidence than incorrect choices for all levels of evidence strength in the model described above; and (ii) reaction times (RT), which has been found to reflect decision confidence as defined above in empirical work (Sanders et al., 2016; Urai et al., 2017).

When assessing the confidence-dependence of the bias adjustment (i.e. changes in history weights), we restricted the model to the immediately preceding trial (lag 1), at which the bias adjustment was expected to be strongest, but we now estimated the weights separately for each of the different levels of previous motion coherence. This enabled us to control for the trial-to-trial variations of stimulus strength, thus isolating the impact of internal trial-to-trial fluctuations of confidence.

In our analysis of the impact of choice accuracy, separate predictors coded for the choice or stimulus categories for each level of (non-zero) previous motion coherence. Because choice accuracy was undefined at 0% coherence, we estimated a single choice weight for previous trials where no decision-relevant sensory evidence was presented. Specifically, we included six regressors in the model that each coded for the previous choice at a given coherence level (zero elsewhere) and we included five regressors that each coded for the previous stimulus category at a given non-zero coherence level (zero elsewhere). To assess the impact of choice accuracy, the stimulus and choice weights were transformed into weights for correct and incorrect choices by re-combining the stimulus and choice weights as described in the section *Modeling the contributions of past correct (and incorrect) choices to current bias* above.

To assess the effect of RTs, we first normalized RT to make it scale positively with confidence, because of its negative scaling with decision confidence (the shortest RTs correspond to the most confident trials (Sanders et al., 2016; Urai et al., 2017): For each observer and condition, we transformed single-trial RTs as follows:

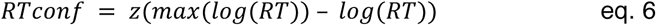

where *z* denoted z-scoring per individual and condition. This transformation was only applied to simplify the interpretation of the corresponding history terms in terms of confidence-weighting. Without this transformation, the resulting weights were qualitatively identical but sign-flipped, thus reflecting the complement to confidence, decision uncertainty (data not shown).

We added a modulation by the above-defined *RTconf* variable to the logistic regression model, as introduced in (Urai et al., 2017). To this end, we added a term describing the interaction between choice and stimulus category at lag 1 with the previous trial’s *RTconf* separately for each previous coherence level: 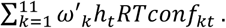 Specifically, the interaction terms in this model were six regressors for previous choice multiplied by previous *RTconf* (one for each coherence level) and five regressors for previous stimulus category multiplied by previous *RTconf* (one for each non-zero coherence level), and a nuisance covariate 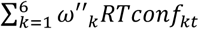 for the main effect of *RTconf.* The full bias term in this model was as follows:

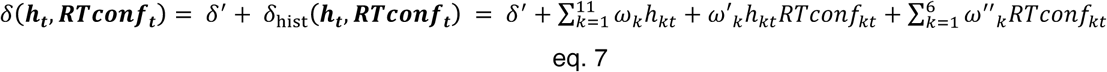

#### Modeling history contributions in synthetic, non-adjusting observers

We performed two sets of simulations to ensure that the context-dependent shifts in the history weights exhibited by participants in Experiment 2 were not just passively ‘inherited from’ the correlated stimulus sequences in the Repetitive and Alternating conditions. The rationale of these simulations was to fit the behavior of synthetic observers. These were matched to the behavior of each of our participants in all parameters, except for the history weights displayed in the biased conditions.

In the first set of simulations, we constructed observers, who based their decisions only on the current stimulus. For each participant, we estimated the parameters of the psychometric function described by eqs. 3 and 4 from the data of the respective biased conditions. This ‘memory-less observer model’ was the set of fitted parameters, but with history bias *δ*_hist_(***h_t_***) set to zero; it allowed us to compute the probability of making an ‘up’-choice (*r_t_ =* 1) for any given stimulus intensity in the absence of any influence of past events. To simulate the model’s performance in the two biased experimental conditions (Alternating and Repetitive), we used the original sequences of stimuli (motion coherences times directions) seen by each observer in these two conditions, and computed the choice probability for each trial by putting the model parameters and stimulus categories in eq.2. Based on these choice probabilities, we then drew the choices on each trial by a weighted coin-flip, resulting in a sequence of choices generated by the model. We then fitted this choice and stimulus sequence, again using the model specified in eqs. 3 and 4 allowing us to estimate the synthetic observers’ stimulus and choice weights. The resulting values served as a reference for the history biases expected as a result of discriminating a biased sequence of stimuli without memory.

The second set of simulations was as the first set of simulations, with the exception that the synthetic observers had the same (non-zero) history weights estimated for the real participants in the Neutral condition, but did not adjust these biases to the biased environments. Again, this enabled us to simulate choice patterns of the synthetic observers exposed to the stimulus sequences used in our actual experiment, and to use these choice patterns to estimate the simulated observers’ choice and stimulus weights, as described for the first set.

In both sets of simulations, we presented the same stimulus sequence 50 times, to average out the effect of binomial noise that was needed to generate choices from the logistic function. This yielded more precise estimates of the model parameters than was possible in the human observers.

### Statistical tests

We used parametric 2-tailed t-tests for all statistical comparisons of regression weights reported in this paper. The rationale was that we could then also provide Bayes factors (Bf), in order to quantify the posterior belief in the null hypothesis given the evidence (Rouder et al., 2009). Bf_10_ < 1/3 indicates evidence in favor of the null hypothesis, Bf_10_ > 3 indicates evidence for the alternative hypothesis, and Bf_10_ = 1 indicates inconclusive evidence. When performing multiple t-tests of regression weights (e.g., across seven lags or coherence levels), false discovery rate correction (Benjamini and Hochberg, 1995) was applied to correct for multiple comparisons.

When testing correlation coefficients computed for individual participants (the so-called ‘adaptivity indices’ defined in Results) against zero, we first Fisher z-transformed the Pearson correlation coefficients and then submitted them to simple t-tests. We used the parametric 2-tailed Steiger’s test (Steiger, 1980) for comparing across-subjects correlations between individual adaptivitiy indices (correlation coefficients) and their proportion of correct choices with the corresponding correlations between individual perceptual sensitivity and the proportion correct choices.

Finally, we used circular statistics, specifically Rayleigh’s test, to assess the clustering of orientations of the lines connecting the weights from Neutral with those from either Repetitive or Alternating conditions, respectively. A Hotelling test (van den Brink et al., 2014) was used to assess the difference in mean directions of adjustment between these two conditions.

The results from all the all regression weights and individual adaptivity indices (correlation coefficients) in Experiment 2 were analogous when replacing the parametric tests with non-parametric permutation tests (Efron and Tibshirani, 1998) with N = 10.000 permutations.

## Results

We here report results from two experiments (referred to as Experiment 1 and 2) quantifying choice history biases during the random dot motion discrimination task that is widely used in neurophysiological studies of perceptual decision-making (Gold and Shadlen, 2007; Siegel et al., 2011; Kelly and O’Connell, 2015). The two experiments aimed to manipulate different aspects of choice behavior. Analyses of behavior from Experiment 1 were previously published (Tsetsos et al., 2015), but those analyses did not assess sequential effects. Here, we re-analyzed these data to quantify the dependence of choice on previous stimuli, choices, and motor responses.

Figure 1C and D illustrates behavioral patterns generated by choice history biases in example observers from the Neutral condition of Experiment 2 (i.e. no correlations among successive stimuli). Figure 1C shows, for one observer, a ‘streak’ of eight repeats of the same choice, followed by five repeats of the other choice. These streaks occur in the face of trial-to-trial variations of the direction of the random stimuli. Critically, such apparent biases towards one or the other choice emerge only locally in time. Choice history biases are therefore distinct from the ‘global’ biases towards one particular choice that result from uneven probabilities of the two stimulus categories or uneven payoffs for the two options (Bogacz et al., 2006; Mulder et al., 2012; de Lange et al., 2013). One way to isolate choice history biases is to fit, for each observer, two separate psychometric functions (relating signed stimulus strength to choice probability), each conditioned on the choice the observer made on the previous trial. Choice history biases are then evident as horizontal shifts between these two functions. Figure 1D displays the resulting functions of three example observers (Neutral condition from Experiment 2) with an intrinsic bias to repeat (left panel, same observer as in Figure 1C) or to alternate choices (right), or no bias (middle). A more comprehensive approach is to explicitly model the relative contribution of previous choices, or other experimental variables from previous trials, to current choice bias (Busse et al., 2011; Fründ et al., 2014) (see Materials and Methods, section *Modeling choice history bias).* We used this statistical modeling approach throughout this paper.

Our analyses pursued two main objectives. First, we aimed to disentangle and compare the contribution of decisional and motor processing stages to the history biases. Second, we aimed to quantify the adjustment of choice history biases to the environment, as a function of varying levels of decision confidence, in the absence of external feedback.

### Experiment 1: Disentangling the impact of previous stimuli, choices, and motor responses

In laboratory tasks, perceptual choice and motor response used for reporting the choice are typically coupled, but can be decoupled with little effect on performance on the current trial (Tsetsos et al., 2015). While there is evidence for either decisional or motor origin of history biases (Akaishi et al., 2014; Pape and Siegel, 2016; St. John-Saaltink et al., 2016), their relative contributions have not yet been systematically compared across several trials in the past. To do so, we reanalyzed data from a previously published study (Tsetsos et al., 2015), in which observers performed a random dot motion task under trial-to-trial variations in the mapping between choice and motor response. The direction of motion was chosen randomly and independently on each trial, so that maximizing performance required basing choices solely on the current stimulus and not on its history (i.e., previous stimuli, choices, or motor responses).

Observers showed a significant tendency to repeat their previous choices (indicated by positive choice weights), but not their motor responses (Figure 2A). The effect of the previous choice on current choice was positive and stronger than the effect of the previous motor response (Figure 2A). The response weights did not differ significantly from zero for any lag (Bf_10_ < 0.45 for all lags). Preceding stimulus categories (up/down) exhibited negative, albeit not statistically significant, weights at longer lags (Figure 2B), possibly reflecting the impact of long-lasting, repulsive effects of direction-selective sensory adaptation mechanisms (Kohn, 2007) on choice behavior.

**Figure 2.**
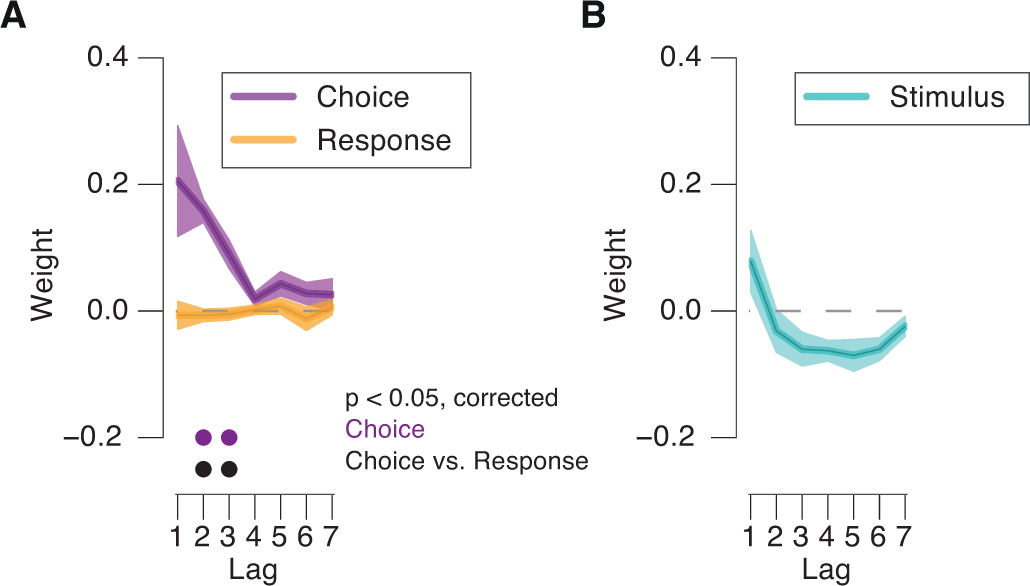
Stronger impact of previous choice than of previous motor response on current bias. **(A)** Impact of previous choices and motor responses as a function of lag. **(B)** As (A), but for impact of previous stimulus categories. Shaded areas, s.e.m.; dots, p < 0.05 (FDR-corrected t-test) across participants.

These results indicate that the commonly observed choice repetition biases are specifically due to previous choices and not the motor responses used to the report them, which has implications for their neural bases (see Discussion). We next investigated the adjustment of choice history biases (under fixed mapping) to varying environmental statistics in order to gain deeper insights into their functional origin and adaptive utility.

### Experiment 2: Confidence-dependent adjustment of choice history biases to the environment

In laboratory tasks used to study perceptual choice, it is common to generate random sequences of the two alternative stimulus categories. But the states of natural environments, and hence the sensory signals generated by them, often exhibit significant auto-correlations across time, so that it might be beneficial for decision-makers to adjust their choice history biases to this correlation structure (Yu and Cohen, 2009). In Experiment 2, we tested for such adjustments, by systematically manipulating the repetition probabilities between the two possible motion directions across three conditions blocked by experimental session: Repetitive, Alternating, and Neutral (two sessions per condition; see Figure 1B and Materials and Methods). Importantly, observers received no external feedback about the correctness of their choices. This enabled us to study the impact of their decision confidence on the adjustment of their choice history biases to environmental statistics.

#### Adjustment of choice history biases to environmental statistics

The manipulation of the environmental statistics had robust effects on observers’ history biases. We visualized those in two complementary ways focusing on different aspects of the data. Both our approaches were guided by the statistical structure of the Repetitive and Alternating conditions, which yielded characteristic profiles of the probability of stimulus repetitions as a function of lag: For both Repetitive and Alternating conditions, repetition probability was most strongly biased (i.e., different from 0.5) at lag 1, and progressively approaching 0.5 for larger lags (Figure 3A). Thus, the strongest effects were expected for events from the preceding trial (i.e., lag 1).

**Figure 3.**
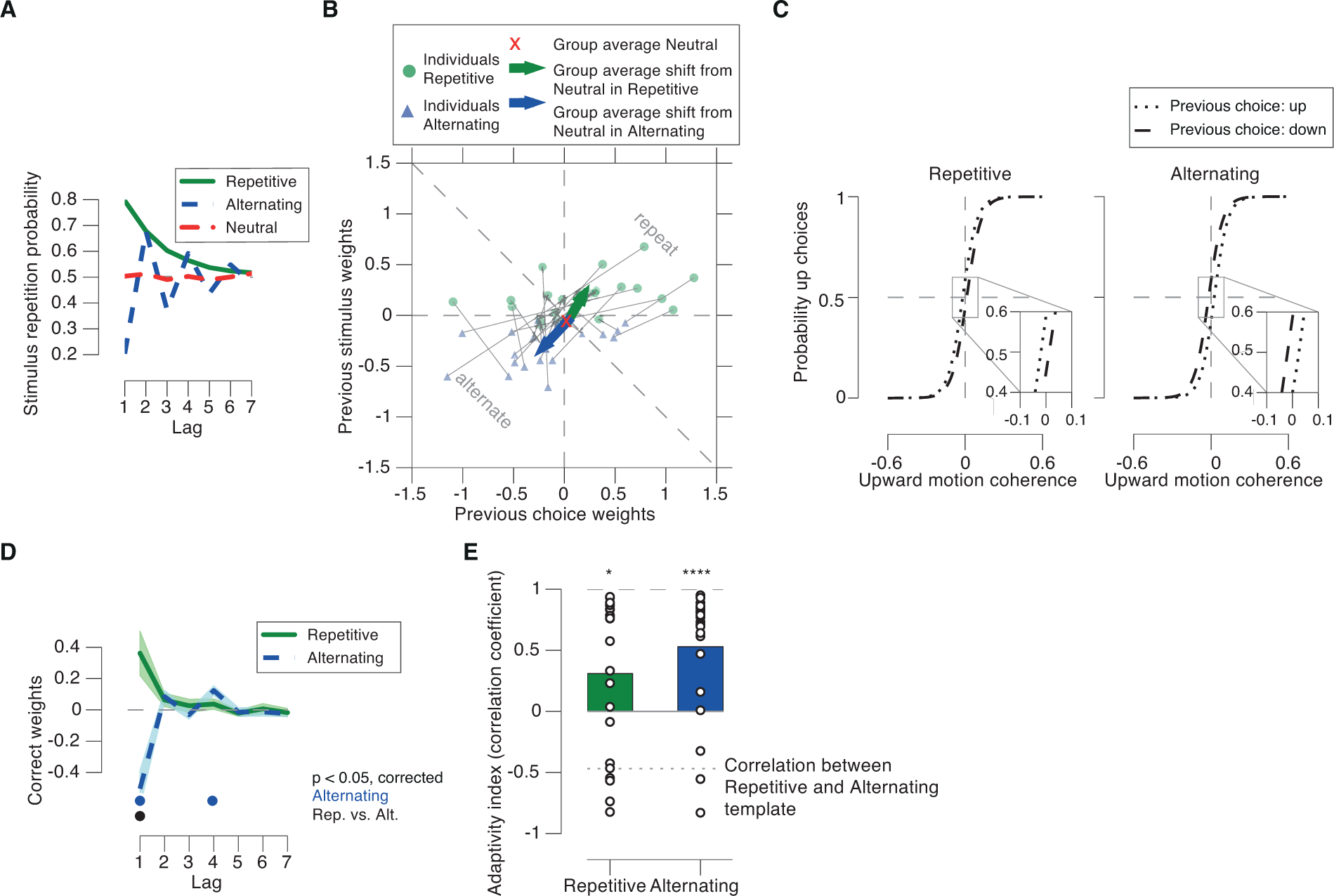
Adjustment of choice history biases to environmental statistics. **(A)** Stimulus repetition probabilities for Repetitive, Alternating and Neutral. In Repetitive, repetition of the motion direction from two trials back could occur due to a sequence of two repetitions or two alternations (probability: 0.8×0.8 + 0.2×0.2 = 0.68). In Alternating, the probability of repetition of the same direction oscillated around 0.5 as a function of lags, with decreasing deviation from 0.5. **(B)** Impact of previous stimuli and choices on current choice for lag 1. Dots, single observers, arrows changes of group mean weights from Neutral (red cross) during Repetitive and Alternating, respectively. **(C)** Psychometric functions conditioned on previous choice (group average). Left: Repetitive, leftward shift from dashed to dotted line corresponding to a bias to repeat the previous choice. Right: Alternating, leftward shift from dotted to dashed line, indicating a bias to alternate the previous choice. **(D)** Correct weights as functions of lags in Repetitive and Alternating. **(E)** Adaptitivity indices (correlation coefficient with the history template) computed from correct kernels from the Repetitive and Alternating conditions. Dotted line, correlation between history templates for Repetitive and Alternating. Shaded areas, s.e.m.; dots, p < 0.05 (FDR-corrected t-test) across participants; *, p < 0.05; **** p < 0.0001.

Our first approach, therefore, focused on the weights for lag 1. When plotting the choice weights against stimulus weights, data points located in the upper-right triangular part indicated a tendency to repeat the previous choice or stimulus categories (up/down), whereas data points in the lower-left triangular part indicated a tendency to alternate (Figure 3B, dashed diagonal line). If observers adjusted their choice patterns to the Repetitive and Alternating environments, their history weights should have shifted in the corresponding directions. This is what we observed (Figure 3B; compare dots of different colors). The weights were close to zero in the Neutral condition (group average, red ‘x’); weights shifted towards repetition in the Repetitive condition (group average, green arrow in Figure 3B), and alternation in the Alternating condition (group average, blue arrow in Figure 3B), respectively. The vector angles of the shift from Neutral were significantly different from uniform (z = 7.69, p = 0.0003 in Repetitive and z = 8.64, p < 0.0001 in the Alternating condition, Rayleigh’s test), and the shift angles were significantly different between Repetitive and Alternating (F(2, 20) = 60.28, p < 0.0001, Hotelling test). The adjustment of choice history bias was also evident when fitting the psychometric function conditioned on the previous choice (as in Figure 1D). Both conditions were characterized by a history-dependent shift, in opposite directions (Figure 3C; difference in shift between Repetitive and Alternating: t(21) = 3.21, p = 0.0042). By contrast, previous choice had no effect on the slope of the psychometric function (difference in history-dependent change in the slope between Repetitive and Alternating: t(21) = −0.60, p = 0.5532, Bf_10_ = 0.2627).

It is noteworthy that the direction of the shift of history weights between the different environmental statistics was largely along the positive diagonal (Figure 3B) that corresponds to equal weights for previous stimuli and choices. Thus, the bias adjustment was largely driven by correct choices (where previous choices and stimuli were identical). We thus used the weight of previous correct choices (i.e., the sum of choice and stimulus weights, see Materials and Methods) in all subsequent analyses as a single metric of the bias adjustment. The importance of previous correct choice for the bias adjustment was indicative of the role of decision confidence, an aspect that we elaborate on in the section *Modulation of choice history bias adjustment by decision confidence* below. AB C

Our second approach focused on an assessment of the full time courses of the history weights. The temporal profiles of the stimulus repetition probabilities in the two biased conditions exhibited markedly different patterns: In the Repetitive condition the temporal profile exhibited a monotonic decay towards 0.5, whereas it exhibited a damped oscillation around 0.5 in the Alternating condition (Figure 3A). The correlation between both time courses for Repetitive and Alternating conditions was negative. In what follows, we refer to these time courses as ‘history templates’, to indicate that these characterize the statistical structure of the environment.

Indeed, participants’ history weights exhibited profiles that were similar to those of the history templates (Figure 3D, compare with Figure 3A). We use the term ‘history kernel’ to refer to the individual courses of the weights for correct choices as a function of lag. We quantified their similarity with the corresponding history templates by means of temporal correlation (Figure 3E). These correlations were significant in both conditions (Repetitive: t(21) = 2.57, p = 0.0179, Alternating: t(21) = 4.83, p < 0.0001). Thus, participants adjusted their history biases to the statistical structure of their environments with a time course matched to the full environmental statistics. In what follows, we refer to this similarity metric as ‘adaptivity index’.

One concern might be that even an observer who only discriminates the current sensory evidence, without any active accumulation of past experimental events, might exhibit similar shifts in the history weights between the Alternating and Repetitive conditions, by virtue of the stimulus statistics propagating into the history weights without any active adjustment of the observer. To address this concern, we simulated the performance of two types of synthetic observers, which were constructed individually for each of our participants. These had the same perceptual sensitivity as each participant, but without any adjustment of stimulus and choice weights to the different environmental conditions (see Materials and Methods). The first set of synthetic observers had stimulus and choice weights of zero. The second set of synthetic observers had the same history biases (i.e., non-zero choice and stimulus weights) as our participants in the Neutral condition, but did not adjust these to the changing environmental statistics. The choice and stimulus weights obtained for both these models were not significantly different from zero (Figure 4A and B) (all p-values > 0.567 and Bf_10_ ranging from 0.22 to 0.62).

**Figure 4.**
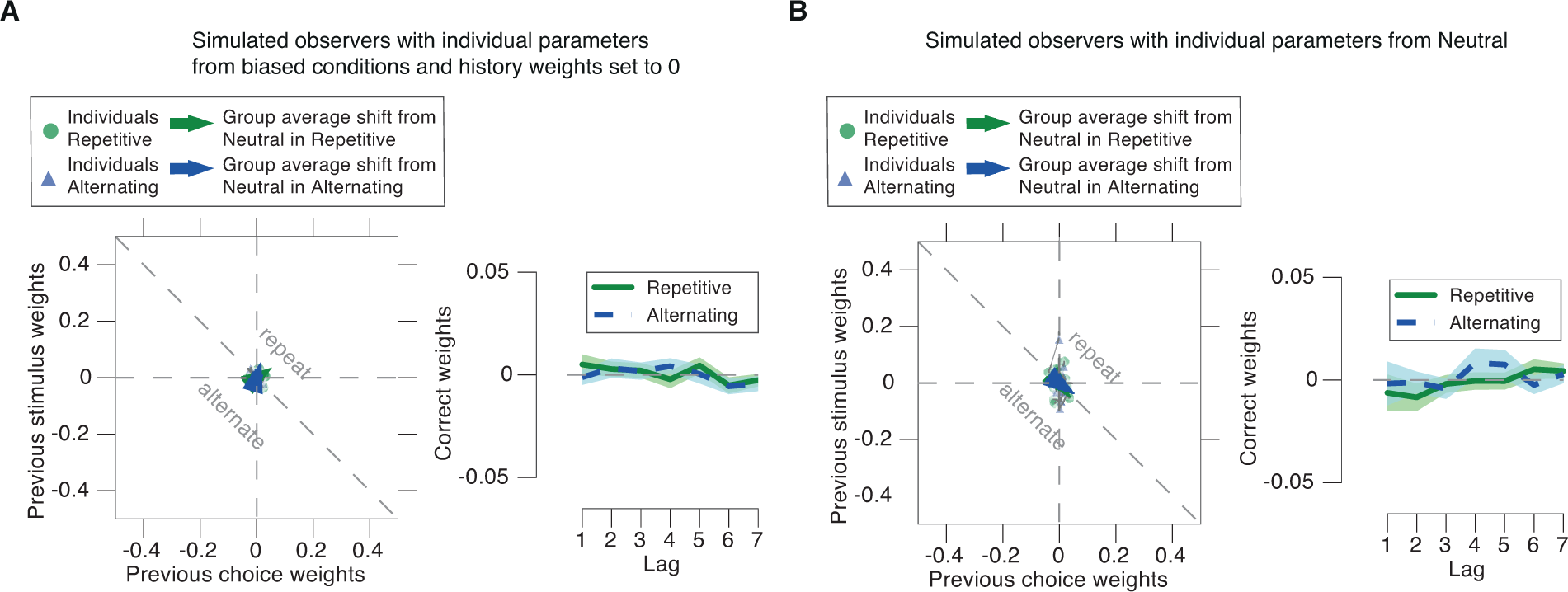
No differences in history weights for synthetic observers without bias adjustment. Results of two simulations of synthetic observers without bias adjustment analyzed as real observer data for Figure 3B, D. (**A)** Synthetic observers with all parameters taken from the real observers’ estimates for Repetitive and Alternating, but history weights set to 0. (**B**) Synthetic observers with all parameters taken from the real observers’ estimates for Neutral. Note the difference in the y-axis scale between these simulated observers and the real observer data in Figure 3. Shaded areas, s.e.m. across synthetic observers.

These simulation results indicate that the effect of the correlations between stimuli on choice patterns was reliably soaked up by the stimulus-dependent part of our statistical model (i.e. the slope of the psychometric function). In other words, the systematic deviations of the history weights between Repetitive and Alternating conditions evident in the real observers were not just passively ‘inherited from’ the correlations evident in the stimulus sequences, but rather due to an active adjustment of their biases.

#### History bias adjustment predicts performance in biased environments

While the bias adjustment was highly consistent across participants, individuals differed in the extent to which they shifted their history biases between conditions (i.e., the magnitude of their adaptivity indices, Figure 3E). We correlated the individual adaptivity indices with the proportion of correct choices to assess their predictive value for overall task performance (Figure 5A). The more strongly observers adjusted, the more successful they were in both, the Repetitive (Figure 5A, left panel) and Alternating (middle) condition. We found no evidence for such an effect in the Neutral condition (Figure 5A, right, Bf_10_ = 0.4276). As expected, perceptual sensitivity (i.e., the slope of the psychometric function) was also strongly predictive of individual performance in all three conditions (Figure 5B). In both biased environments the adaptivity index was similarly predictive of performance as perceptual sensitivity (Repetitive: Steiger’s test z = 0.28, p = 0.7791; Alternating: z = 0.4, p = 0.6893), while sensitivity was a better predictor in the Neutral environment (z = 2.70, p = 0.0068). In other words, the adjustment to the environmental statistic can be about as important for maximizing reward rate as sensitivity to the momentary sensory evidence.

**Figure 5.**
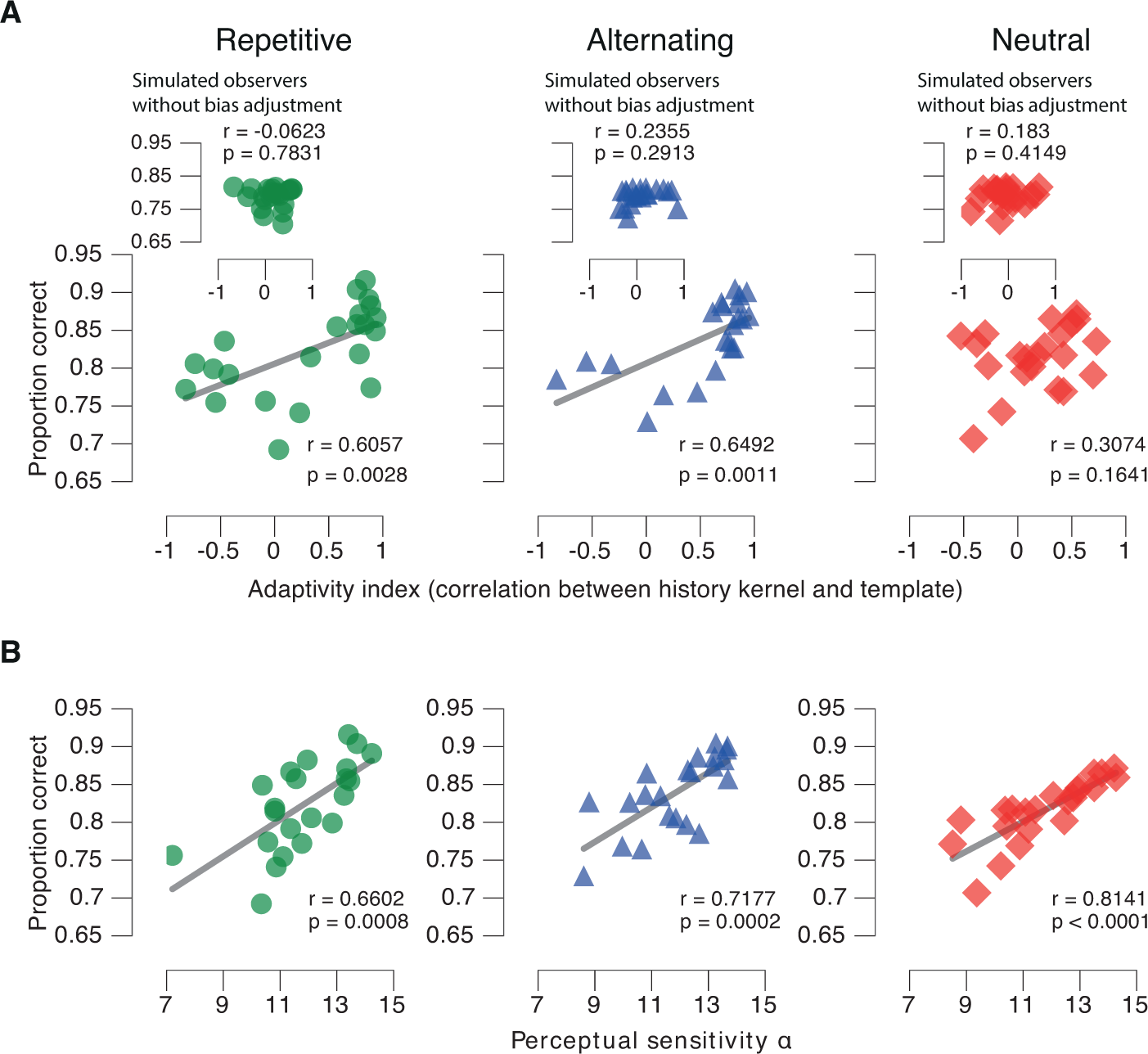
Behavioral performance depends on bias adjustment and perceptual sensitivity. **(A)** Correlation between adaptivity index and proportion of correct choices. Left: Repetitive. Middle: Alternating. Right: Neutral. Insets: Correlations for the simulated observers without bias adjustment (compare Figure 4A). **(B)** Correlation between sensitivity (i.e. slope of the psychometric function) and the proportion of correct choices. Left: Repetitive. Middle: Alternating. Right: Neutral.

Taken together, our results reported so far supported the idea that participants accumulated internal signals from their previous correct decisions into biases for their current choice – a process that adjusted their behavior to the statistics of their environment and improved performance. Some previous accounts of sequential effects have postulated the accumulation of external variables, such as stimulus repetitions (Yu and Cohen, 2009; Meyniel et al., 2016), performance feedback (Abrahamyan et al., 2016), or reward (Sugrue, 2004). Our experimental conditions precluded any of the above: Observers performed under generally high uncertainty about the veridical stimulus identities, and they did not receive external feedback about choice outcomes. We reasoned that, under these conditions, observers may have accumulated the internal decision variables, on which they based their choices in a context-dependent manner (i.e., with opposite sign for Repetitive and Alternating environments). This interpretation is in line with a normative model of sequential effects (Glaze et al, 2015). In statistical decision theory, as well as in neural signals observed in the brain, the decision variable not only encodes the categorical choice, but also the graded confidence about that choice (Kepecs et al., 2008; Kiani and Shadlen, 2009; Hebart et al., 2016). Consequently, we reasoned that the impact of previous choices on current bias should depend on the confidence associated with the previous choices. Our final set of analyses tested this hypothesis.

#### Modulation of choice history bias adjustment by confidence

We here use the term ‘decision confidence’ in a statistical sense, to refer to the posterior probability that a choice is correct, given the evidence (Kepecs et al., 2008; Pouget et al., 2016; Sanders et al., 2016; Urai et al., 2017). The key features of a model formalizing this construct are reproduced in Figure 6A (Kepecs et al., 2008; Sanders et al., 2016; Urai et al., 2017; see Methods). This definition of confidence is agnostic about the link to the subjective sense of confidence, or the ability to report this sense of confidence (but see Sanders et al, 2016).

**Figure 6.**
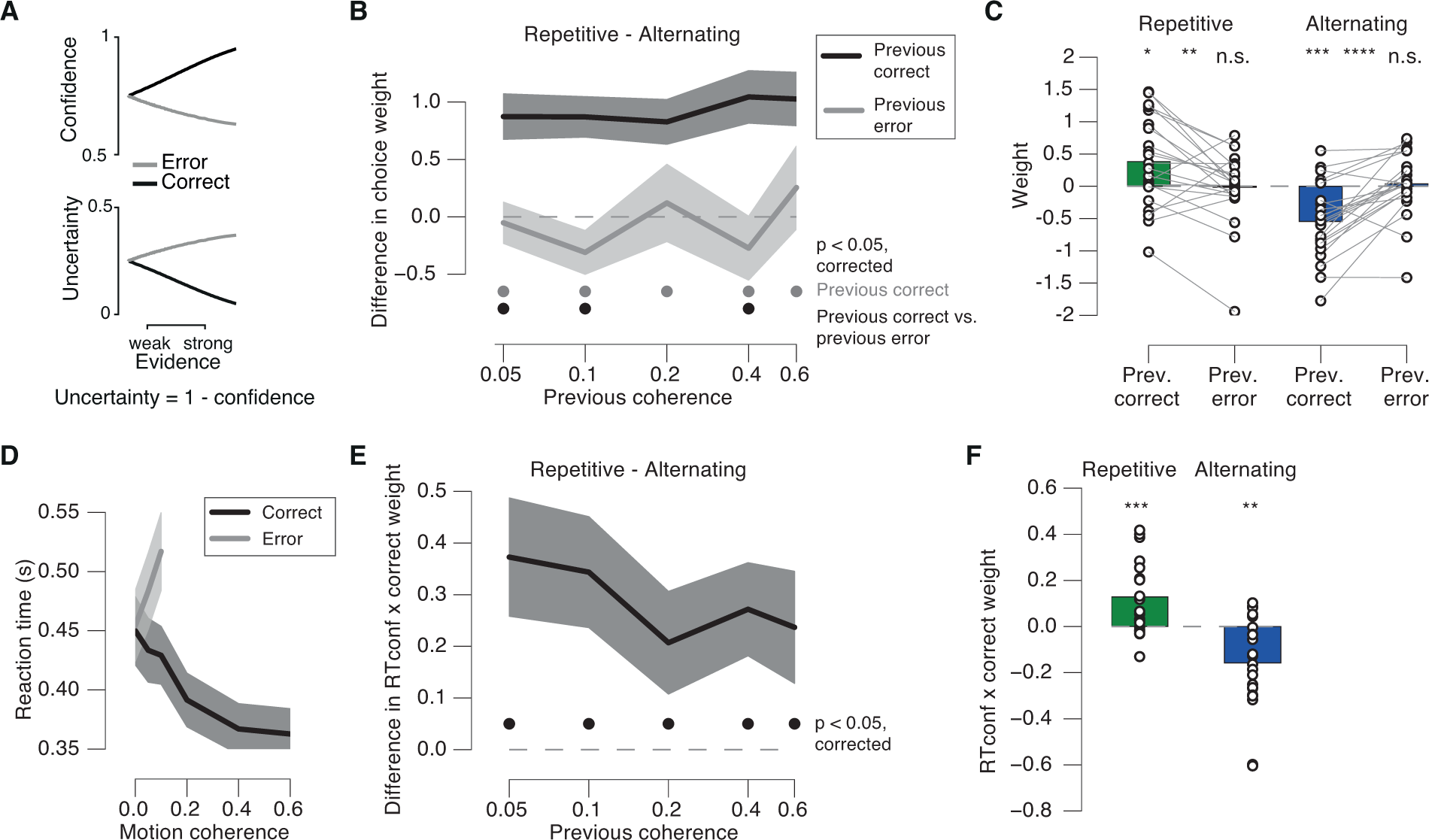
Modulation of bias adjustment by proxies of confidence in previous choice. **(A**) Scaling of model-based confidence and uncertainty with evidence strength on correct and error trials. Adapted from (Urai et al., 2017) under a CC-BY 4.0 license. **(B)** Difference between previous choice weights from Repetitive and Alternating, sorted by previous choice correctness and coherence. **(C)** Comparison between previous correct and incorrect weights, for Repetitive and Alternating. Weights were first calculated separately for each previous coherence level and then pooled across coherence. **(D)** Reaction time as function of motion coherence sorted by correctness (pooled across Repetitive and Alternating). **(E)** Difference between previous *RTconf x correct* modulation weights from Repetitive and Alternating, sorted by previous coherence. See main text for details of the multiplicative modulation model. **(F)** *RTconf x correct* weights for Repetitive and Alternating. Modulation weights were first calculated separately for each previous coherence level and then pooled across coherence. Shaded areas, s.e.m.; dots, p < 0.05 (FDR-corrected t-test) across participants; *, p < 0.05; **, p < 0.01; ***, p < 0.001, ****, p < 0.0001.

We used two experimental variables consistent with this definition of confidence: accuracy and RT. Correct choices are overall associated with higher confidence for all non-zero evidence strengths (i.e., coherence levels, Figure 6A, top). The scaling of RT with motion coherence exhibited the same characteristic signature as uncertainty (i.e., the complement of confidence) as reported in previous studies (Sanders et al., 2016; Urai et al., 2017): RT decreased with coherence for correct choices, but increased for incorrect choices (Figure 6D, compare to Figure 6A). Linear regression revealed an opposite-signed relationship between motion coherence and RT, separately for correct (β = −0.150, s.e.m. = 0.027, p = 0.005) and error trials (β = 0.628, s.e.m. = 0.025, p = 0.025).

As predicted, the leverage of the preceding choice on bias adjustment (i.e., difference between Repetitive and Alternating) was larger when the previous choice was correct than incorrect, even when controlling for the level of previous motion coherence (Figure 6B, C). There was a significant effect of previous correct choice at all of the previous coherence levels, while there was no such effect for previous incorrect choices at any of the previous coherence levels (Figure 6B; Bf_10_: 0.23, 0.69, 0.23, 0.34, and 0.28 for previous coherence levels: 0.05, 0.1, 0.2, 0.4, and 0.6, respectively).

When pooled across previous coherence levels, the weights for previous correct choices deviated significantly from zero in both conditions (Figure 6C; Repetitive: t(21) = 2.58, p = 0.0174; Alternating: t(21) = −4.42, p = 0.0002). Again there was no such effect for previous errors (Figure 6C; Repetitive: t(21) = −0.13, p = 0.8985, Bf10 = 0.22; Alternating: t(21) = 0.35, p = 0.7309, Bf10 = 0.24). The weights were significantly larger for correct than incorrect previous choices in Repetitive (t(21) = 3.06, p = 0.0060, and the other way around in the Alternating (t(21) = −5.19, p < 0.0001 Figure 6C). Please note that while the weights were averaged across previous coherence levels for visualization in Figure 6C, they were first estimated separately for each previous coherence in order to factor out effects of trial-to-trial fluctuations in sensory evidence strength (see Materials and Methods). These results were qualitatively identical after random subsampling of the correct trials, so as to match the smaller number of incorrect trials for each previous coherence level (data not shown), ruling out the concern that the stronger bias adjustment after correct choices may have been due to the larger number of correct than error trials. In sum, these results were consistent with the idea that the weight of choices in the across-trial accumulation process depended on internal (i.e., stimulus-independent) fluctuations in decision confidence.

To assess the modulatory effect of the second confidence proxy, RT, on the bias adjustment, we built on an extension of the statistical model by multiplicative interaction terms. This quantified the degree to which the impact of previous correct choices on current bias was modulated by previous RT (see Materials and Methods for details). In these model fits, we transformed RT to scale positively with decision confidence, a variable we refer to as *RTconf* (Materials and Methods). Again, we split trials by their motion coherence to assess the modulatory effect of *RTconf* on the impact of correct choices (i.e. the weights for the interaction term *RTconf x correct)*, over and above variations in evidence strength.

Larger values of *RTconf* were associated with a stronger impact of the previous (correct) choice on the current bias (Figure 6E, F), an effect that was robust even when we evaluated each previous coherence level separately (Figure 6E). When pooled across previous coherence levels, the interaction weight was significantly larger than zero in the Repetitive condition (Figure 6F; t(21) = 3.84, p = 0.0009), indicating a confidence-dependent enhancement of the tendency to repeat correct choices in that condition. Conversely, the interaction weight was significantly smaller than zero in the Alternating condition (Figure 6F; t(21) = −3.71, p = 0.0013), indicating a confidence-dependent enhancement of the tendency to alternate correct choices. Thus, even within the correct choices, evidence-independent fluctuations in the associated confidence (indexed by *RTconf*) boosted their impact on future choice bias.

Taken together, two independent proxies of decision confidence, choice accuracy and reaction time, both supported the conclusion that decision confidence boosted the adjustment of choice history biases to the structure of the environment.

## Discussion

Choice history biases are a pervasive phenomenon in perceptual decision-making (Fernberger, 1920; Fründ et al., 2014). Here, we showed that these biases were largely dominated by categorical choices rather than motor responses and could be flexibly adjusted to environmental statistics even in the absence of feedback about choice outcome. In line with recent normative accounts, the strength of this adjustment was modulated by previous decision confidence. In environments with strong sequential structure it governed individual performance to a similar extent as perceptual sensitivity. Taken together, our results yield new insights into the functional origins and adaptive utility of choice history biases, with direct implications for their neural bases.

An important novel contribution of our study is the discovery of a confidence-weighted adjustment of choice history biases to changing environments. We propose that this was due to a context-dependent accumulation of decision variables across trials. A similar accumulation process has been proposed to explain sequential effects under strong and unambiguous evidence (Yu and Cohen, 2009; Meyniel et al., 2016). Consequently, those latter models describe the accumulation of external observables rather than internal decision variables. The latter are often dissociated from external observables when the decision-maker is uncertain about the environmental state due to degraded evidence. While temporal accumulation is a widely established mechanism in perceptual choice (Bogacz et al., 2006; Gold and Shadlen, 2007; Ratcliff and McKoon, 2008; Wang, 2008; Ossmy et al., 2013), previous models focus on the within-trial accumulation of the momentary sensory evidence. Across-trial accumulation of information is long established in the theory of reinforcement learning, but there it pertains to the accumulation of rewards (i.e., external signals about choice outcome) and spans substantially longer timescales than sensory evidence accumulation (Sutton and Barto, 1998; Sugrue, 2004; Glimcher, 2011). Our current results are indicative of an accumulation mechanism that operates (i) with a timescale situated in between those established for sensory evidence accumulation and action value learning, and (ii) on internal decision variables, which themselves result from the faster (within-trial) accumulation of sensory evidence.

Such a context-dependent across-trial accumulation of decision variables has been postulated by a recent normative account (Glaze et al., 2015), and shown to account for history biases in simple saccadic choice (Kim et al., 2017). Little is currently known about the neural basis of this process. Our current work sets the stage for probing into its neural basis, by experimentally establishing key behavioral hallmarks of this accumulation process within the most widely used task in the neurophysiology of perceptual decision-making (Gold and Shadlen, 2007; Siegel et al., 2011; Kelly and O’Connell, 2015). It will now be important to explore the underlying mechanisms through direct recordings of neural activity under conditions as used here.

One recent study provided similar evidence for an effective adjustment of human observers to changing environmental statistics (Abrahamyan et al., 2016). Our current results and those from Abrahamyan et al (2016) complement each other in quantifying the adaptability of human choice history biases. In Abrahamyan et al. the nature of the change in the environment was different from the one we have used here: In their study, observers received unambiguous feedback about the outcome of each choice, and the manipulation of the stimulus sequence depended on the participants’ success or failure. Consequently, the process adjusting history biases likely depended on the combination of choices and their outcome. By contrast, in our study, the environments differed in their statistical structure independent of observers’ choices. Furthermore, participants remained uncertain about their choice outcomes, and therefore could only base their history biases on internal signals. This was likely the key aspect that mediated the confidence-weighting of the impact of previous choices on current bias in our present study. Thus, the adjustment effects in our study and the one from Abrahamyan and colleagues (2016) likely resulted from distinct computational mechanisms.

The interpretation provided above as well as the normative framework by Glaze et al (2015) offer a natural interpretation of the modulatory effect of decision confidence on the adjustment of choice behavior. In statistical decision theory, as well as in neural signals observed in the brain, the decision variable does not only encode the categorical choice, but also the graded confidence about that choice (Kepecs et al., 2008; Kiani and Shadlen, 2009; Hebart et al., 2016). This quantity is the best proxy for the true state of the environment available to the decision-maker in the absence of external feedback. A decision variable large in magnitude implies large confidence and predicts accurate as well as fast decisions (Sanders et al., 2016). Thus, across-trial accumulation of decision variables into choice history bias predicts that correct or fast choices have a stronger impact on the history bias adjustment to environmental statistics – just as we observed in our second experiment. The same idea can account for the observation (Urai et al., 2017) that ‘intrinsic’ history biases emerging under random stimulus sequences, regardless of their direction (i.e., towards alternation or repetition), are weaker following low-confidence decisions (i.e., long reaction times). In such contexts, corresponding to the Neutral condition in our Experiment 2, observers’ biases might result from biased internal representations of the environmental structure (i.e., biased ‘subjective hazard rates’ in the model by Glaze et al, 2015)). Taken together, confidence-weighting of the impact of previous choices on current bias may be a diagnostic feature of the accumulation of graded decision variables across trials that we propose as a mechanism underlying the history bias adjustment in our experiment.

In our account the strength of history bias adjustment depends on the magnitude of the decision variable, which is also the sole source of variations in confidence in the confidence model from Figure 6A (Kepecs et al, 2008). According to this model the difference in confidence between correct and error trials increases as a function of stimulus strength. Thus, one would expect the impact of correctness on bias adjustment to also increase as a function of stimulus strength. Such an increase was not evident in our data (Figure 6B). A possible explanation is that choice accuracy was less closely coupled to the decision variable than postulated by the confidence model from Figure 6A. For example, some errors will be caused by noise downstream from the decision variable and consequently not affect the bias adjustment. Thus, motor errors will counteract the dependence of history bias adjustment on correctness. Because misperception of the true stimulus category becomes less likely with stronger evidence, the relative contribution of motor errors to incorrect choices will increases as a function of evidence strength. This might explain why the effect of previous correctness on history bias adjustment did not increase as a function of previous evidence strength in our data. Similar considerations hold for our second confidence proxy, reaction time.

Our analyses revealed that the contributions of previous stimuli, perceptual choices, and motor responses were dissociable in terms of their strength, sign, and time course. Importantly, the dominant and consistent bias in standard conditions with random stimulus sequences, was to repeat preceding choices, rather than motor responses. Two recent studies similarly decoupled perceptual choice and motor response (Akaishi et al., 2014; Pape and Siegel, 2016). One of them (Pape and Siegel, 2016) showed that a bias to alternate response hands from trial to trial systematically contributed to sequential effects, due to activity dynamics within motor cortex. This motor response alternation bias was superimposed onto a choice repetition bias in their study, but Pape and Siegel (2016) did not compare the magnitude and time course of these two effects directly. When performing such a direct comparison, we here found the contribution of previous choices to be significantly stronger, and more prolonged in time. The predominance of choices over motor responses is consistent with the results (focusing on the preceding decision only) from Akaishi et al (2014). Taken together, the data by Akaishi et al (2014) and our present study indicate that history biases in perceptual decision-making are governed by decision variables encoded in an abstract, action-independent format. Such representations of the decision variable exist in associative brain regions, such as posterior parietal or prefrontal cortex (Bennur and Gold, 2011; Hebart et al., 2012, 2016), which also exhibit the short-term memory dynamics necessary for the persistence of biases in the decision-making machinery (Wang, 2002; Bonaiuto et al., 2016; Morcos and Harvey, 2016).

We conclude that human observers accumulate action-independent, graded decision variables across trials towards biases for upcoming choices in a context-dependent manner. This process enables observers to adjust their choice behavior to environmental statistics in the absence of unambiguous information about choice outcome. Our findings are in line with normative theory and constrain the candidate neural sources of choice history biases.

## Acknowledgements

We thank Ainhoa Hermoso, Alexandre Hyafil, Jaime de la Rocha, and Florent Meyniel for comments on the manuscript. This research was supported by: German Research Foundation (DFG), DO 1240/2-1, DO 1240/3-1, and SFB 936 projects A7/Z1 (to T.H.D.); and German Academic Exchange Service (DAAD, to A.E.U.).

